# Deep learning-based pan-cancer classification model reveals cancer-specific gene expression signatures

**DOI:** 10.1101/2021.03.15.435283

**Authors:** Mayur Divate, Aayush Tyagi, Derek J Richard, Prathosh A Prasad, Harsha Gowda, Shivashankar H Nagaraj

## Abstract

The identification of cancer-specific biomarkers and therapeutic targets is one of the primary goals of cancer genomics. Thousands of cancer genomes, exomes, and transcriptomes have been sequenced to date. In this study, we conducted a pan-cancer analysis of transcriptome datasets from 37 cancer types provided by The Cancer Genome Atlas (TCGA) in an effort to identify cancer-specific gene expression signatures. We employed deep neural networks to train a model on the transcriptome profile datasets for all cancer types. The model was validated, and its predictive accuracy was determined using an independent dataset, achieving > 97% prediction accuracy across cancer types. This strongly suggests that there are distinct gene expression signatures associated with various cancer types. We interpreted the model using SHapley Additive exPlanations (SHAP) to identify specific gene signatures that significantly contributed to the classification of cancer types. In addition to known biomarkers, we identified several novel biomarkers in different cancer types. These cancer-specific gene signatures are valuable candidates for future studies of their potential utility as cancer biomarkers and putative therapeutic targets.

## Introduction

The identities and phenotypes of different cell types are governed by underlying gene expression patterns. While all cells contain the same genetic information except activated B and T cells, and red blood cells, only a subset of genes is expressed in a given cell type. The identification of cell/tissue-specific gene expression signatures can highlight the molecular markers that define cell identities. This is particularly useful in oncogenic contexts, as the identification of specific biomarkers and therapeutic targets is important to the effective diagnosis, monitoring, and treatment of cancers. For example, prostate-specific antigen (PSA) levels are elevated in prostate cancer patients[1], and it is thus used as a screening and diagnostic biomarker for prostate cancers[2]. Molecular markers are also useful when predicting patient outcomes. The elucidation of cancer-specific biomarkers has, as a result, been a longstanding goal of gene expression-related research. However, most prior studies have been restricted to one cancer type, with gene expression patterns of cancer tissues being compared to those of adjacent normal tissue. In order to identify cancer-specific biomarkers, pan-cancer studies that explore transcriptomic patterns across different cancer types are warranted. Until recently, gene expression profiles from multiple cancer types were not available. However, the advent of high-throughput sequencing methods has revolutionized cancer genomics and transcriptomics studies. International cancer consortiums including The Cancer Genome Atlas (TCGA) and the International Cancer Genome Consortium (ICGC) have sequenced hundreds of samples from multiple cancer types and made this data available to the scientific community [3, 4]. Systematic analyses of these pan-cancer datasets are now possible, offering an opportunity to clarify cancer-specific gene signatures.

Various algorithms have previously been used for transcriptomic signature profiling, including shrunken centroids[5, 6], penalization[7], clustering[8], differential expression[9, 10], and machine learning approaches [11, 12]. Recent advances in new deep learning computational methods have enabled researchers to utilize multilayer neural networks to analyze these large datasets. Deep learning is a cutting-edge artificial intelligence technique that provides a very powerful framework for supervised learning[13]. In comparison to traditional machine learning methods, deep learning methods are well in function approximation and data representation. According to the universal approximation theorem[14, 15], a feedforward network with linear output and at least one hidden layer with any squashing activation function can approximate any decision boundary to arbitrary closeness if provided with enough hidden units. Importantly, deep learning has already shown higher accuracy and superior performance relative to traditional machine learning techniques, including support vector machines and random forest approaches, and has improved computer vision by making important advances[13]. For instance, recent architectures such as ResNet[16] and EfficientNet[17] achieved ~90% classification accuracy on ImageNet [18] database which consists of 3.2 million images of 1000 different objects. In the healthcare domain, Google developed a deep learning model that predicts lung cancer risk using computed tomography (CT) images [19]. Researchers have applied deep learning not only to medical images but also to various omics data types [20, 21]. For example, some studies have employed deep learning to predict cancer types based upon mutation [22–24] and gene expression data [25–27]. Machine learning and deep learning have also been applied for cancer type classification and gene signatures identification, albeit with moderate levels of accuracy. Examples include a genetic algorithm and k-nearest neighbours (GA/KNN) based machine learning method that identified 20 gene expression signatures with 90% accuracy [28] and a convolutional neural network (CNN) model that classified cancer types with 95% accuracy [27].

Herein, we conducted a comprehensive pan-cancer gene expression analysis using deep learning to analyze TCGA RNA-sequencing data to identify cancer-specific gene signatures. We developed a model that was able to accurately classify all cancer types based on transcriptomic data. Interpretation of this model revealed 976 genes that constitute cancer-specific gene expression signatures, including particular genes that can effectively discriminate among different cancer types. The DNN model developed using these gene signatures was able to classify different cancer types with over 97% accuracy. These gene signatures can be used for the development of more reliable gene panels for cancer diagnosis and monitoring. In addition, this DNN model can be extended to establish the origins of cancers of unknown primary.

## Materials and Methods

### Gene expression data collection and pre-processing

We obtained fragment per million per kilobase (FPKM) values for 14,237 tumor samples representing 39 cancer types from the TCGA data portal (https://portal.gdc.cancer.gov) using the GDC data transfer tool (August 2020). We analysed only protein-coding genes and excluded mitochondrial genes. In order to reduce noise and to focus on genes that were reliably expressed, we excluded genes that did not exhibit expression levels ≥ 5 FPKM in at least 50% of samples in a given cancer type. These gene lists were then merged to create a non-redundant gene list for model construction. FPKM values were log10 transformed before using the data for the model training and testing. Sample labels (i.e. cancer types) were encoded using OneHotEncoder from the sklearn python package. Samples of each cancer type were randomly assigned to three data sets: 1) a training set (70%), 2) a testing set (20%), and 3) a validation set (10%). The training and validation sets were used in the training phase of the model, while the test set was used to evaluate model performance. Samples and associated encoded labels were individually supplied during the training, validation, and testing of the model.

### Unsupervised clustering

T-distributed stochastic neighbour embedding (t-SNE) is an unsupervised non-linear dimensionality reduction technique [29]. It works on the following principle, samples that are closer in higher dimensions should remain close to each other in lower dimensions. It uses t-student distribution with one-degree of freedom to compute the similarity between data points mapped in lower-dimensional space. In this way, while performing dimensionality reduction, t-SNE keeps similar samples nearer and dissimilar ones apart. Moreover, t-SNE can explain the polynomial relationship between features of non-linear data better than principal component analysis (PCA) and is well suited for visualization of high-dimensional datasets. Unlike t-SNE, PCA is greatly affected by outliers. Thus, we employed t-SNE before and after log-transformation of data for data visualization. For this, the t-SNE method from the sklearn package was used for projecting data into 2-D by setting the random_state to 123 and the rest of the parameters at default values.

### Implementation details

We used a deep neural network (DNN) for developing, testing, and validating the model for the classification of cancer types based on gene expression data. This network was implemented using Python3 and TensorFlow 2.0. The DNN consisted of five fully connected (FC) hidden layers with 500, 250, 125, 100, and 75 nodes. The output layer had 37 nodes, each representing one of the cancer types. The rectified linear unit (ReLU) activation function was used for all hidden layers, while softmax was used for the output layer. The model was trained with a learning rate of 0.00001 and Adam optimizer [30]. As it was a multiclass classification problem, we used categorical cross-entropy as loss function and one-hot encoded labels. In order to avoid the model overfitting, the training data, we used early stopping to monitor the loss function with patience equal to 3 and with the maximum number of permitted epochs being 50.

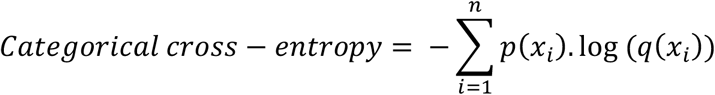

Where *n* is the total of number cancer types, *p(x*_*i*_) is the probability of cancer type *x*_*i*_ and q(*x*_*i*_) is the predicted probability of cancer type *x*_*i*_.

### Evaluation of model performance

It is important to assess model performance using statistical scores in order to establish whether the model requires further improvement. Accuracy is among the most commonly used evaluative metrics, but tends to be biased towards over-represented classes for class imbalanced data. As such cases exist in our dataset, we also used precision, recall, and F1-score values to assess model performance on the test data.

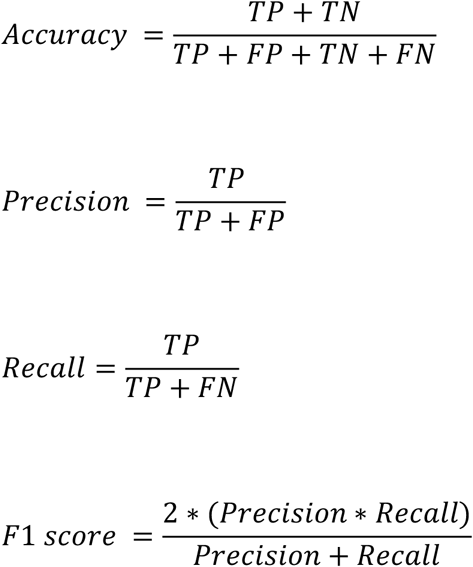

Where TP indicates True Positives, TN indicates True Negatives, FP indicates False Positives, and FN indicates False Negatives. To facilitate model evaluation, we also performed 5-fold cross-validation to ensure the generalizability of the model.

### Model interpretation with SHAP values

To identify the most important model driving features (genes), we conducted model interpretation using SHapley Additive exPlanations (SHAP) values [31]. SHAP values were used to calculate the importance of a given feature by comparing model predictions with and without that feature over all possible combinations of features sampled from the dataset. Since the order in which a model views features can affect the resultant predictions, this computation is repeated in all possible orders to compare the features impartially. A feature that does not affect the predicted value is expected to produce a SHAP value of 0. However, if two features contribute equally to the prediction, then they should have the same SHAP values. Compared to LIME[32] which used local surrogate models to interpret individual predictions, the SHAP method is more accurate and consistent with human intuition as well as has better computational efficiency[26]. We built 10 different DNN models using randomly split TCGA data and SHAP values for each of those models were computed using DeepExplainer, which is based on the combination of SHAP and deepLIFT [33], function from the python SHAP package. Then, the top 20 genes were selected for each cancer type in every model based on absolute mean SHAP values. Then these genes were then compared based on median expression values to establish expression signatures.

## Results

### Unique gene expression signatures are associated with different cancer types

All the cells in the human body share the same genetic information with the exception for life time acquired mutations [34]. Over the course of development, cells differentiate and become specialized to perform specific functions [34]. These changes in cell phenotypes are modulated by changes in gene expression patterns. As such, cell identity and phenotype are ultimately determined based upon these underlying gene expression patterns. This suggests that unique gene expression signatures are associated with each cell type.

To develop the present model, we used transcriptomic data from 14237 samples from 39 cancer types available on the TCGA data portal (Table 1), including adenocarcinomas, squamous cell carcinomas, haematological malignancies, neuronal tumors, melanomas, germ cell tumors, soft tissue tumors, and other cancer types (Table 1). These data were class imbalanced. For example, whereas the breast cancer class had the highest number of samples (1176), only 34 cholangiocarcinoma samples were available (Table 1). We combined proximal cancers, as they show high molecular similarity. For example, colon and rectum data were combined and labeled as colorectal cancer (CORE). Similarly, uterine carcinosarcoma (UCS) was combined with uterine corpus endometrial carcinoma (UCEC). As a result, the data used for this analysis comprised 37 cancer types (Table 1).

**Table 1:**
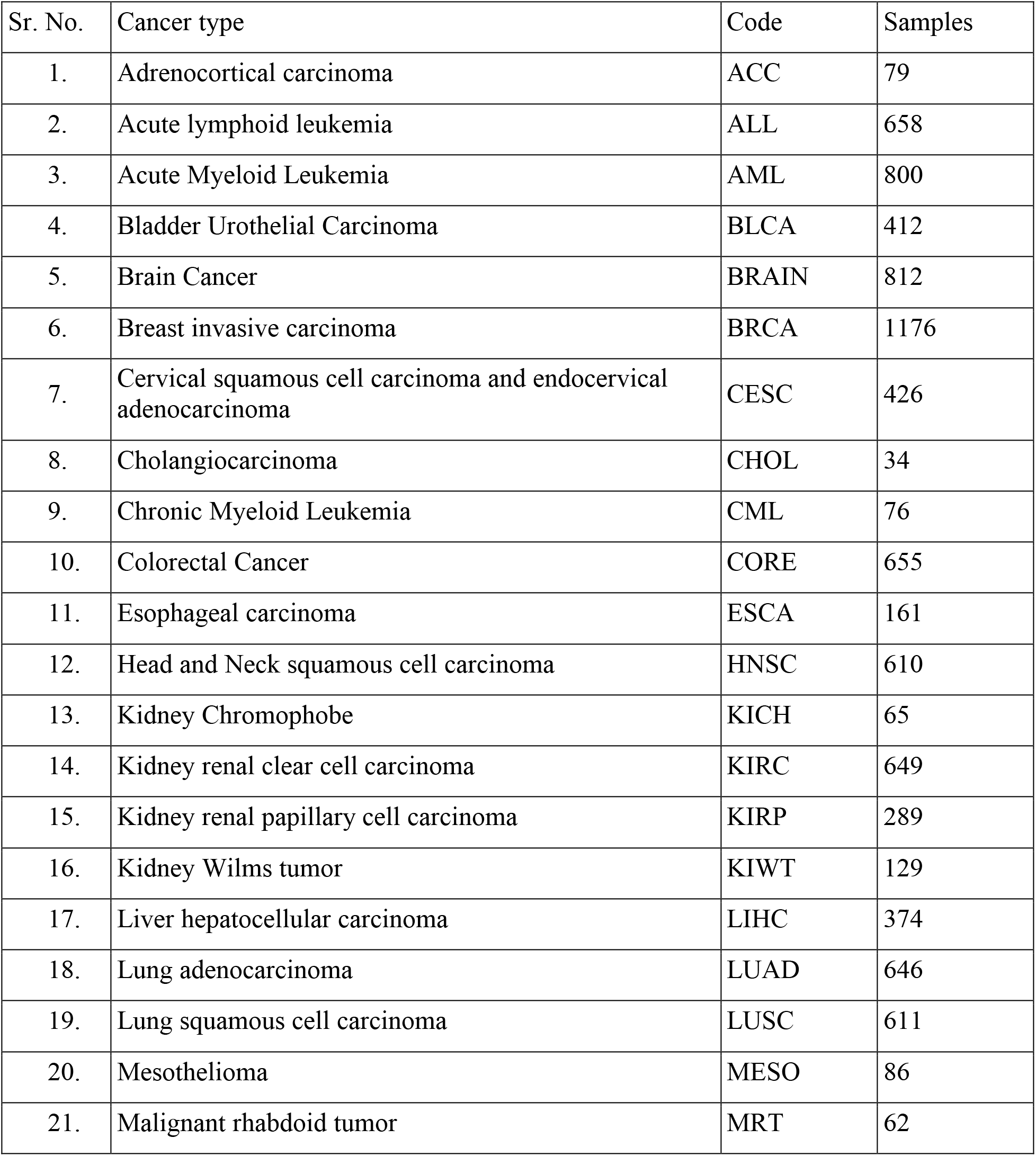
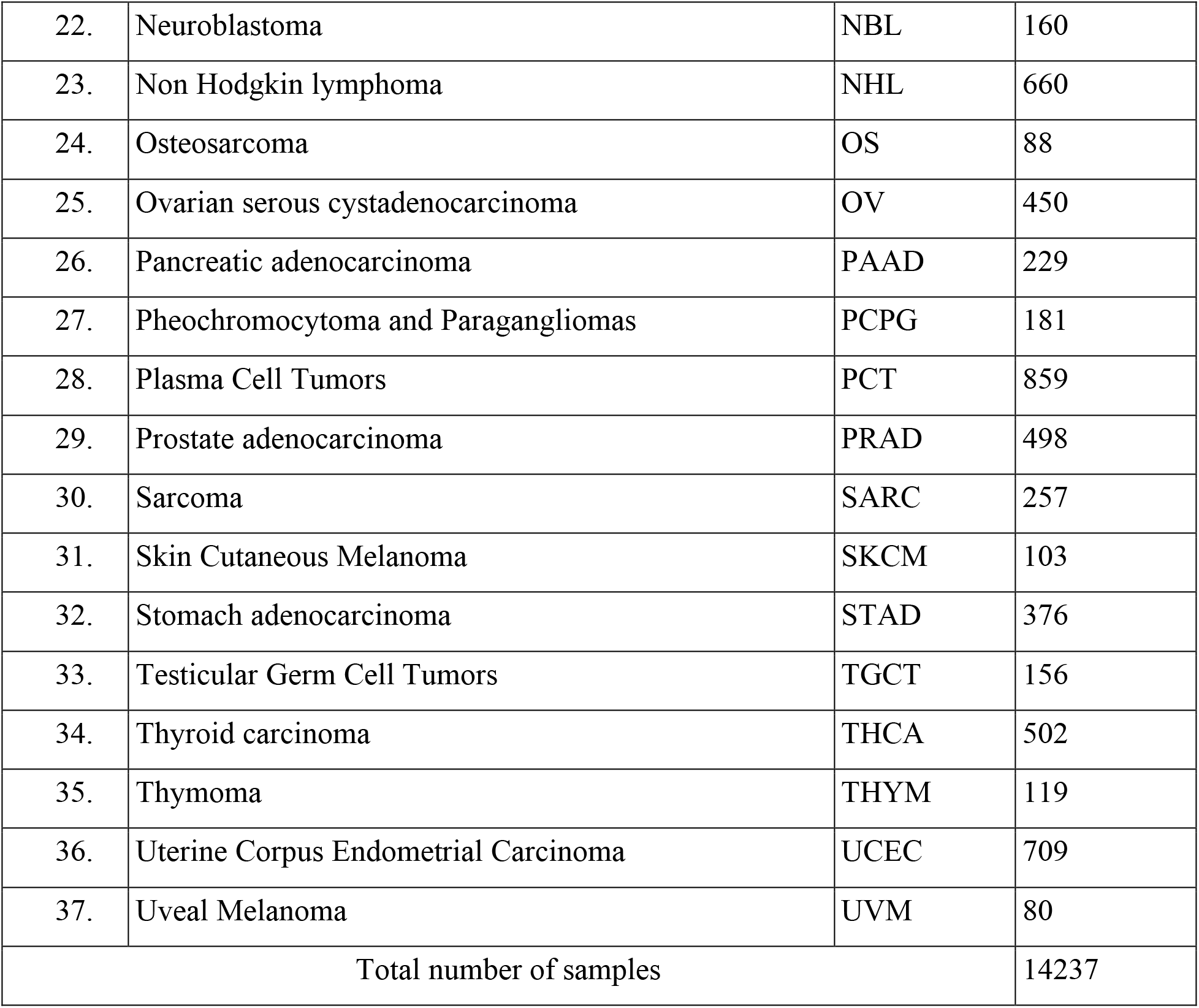
Total numbers of tumor samples and cancer codes of different cancer types used in this study.

Even though RNA-seq is one of the most powerful techniques for assaying gene expression, accurate noise-free quantification remains a challenge [35]. Deep learning models are sensitive to noise in the underlying data, and genes that are expressed at low levels represent a potential source of noise as their measurement across samples can be inconsistent. To eliminate such noise and consider only genes that were reliably expressed in tissues of interest, we filtered data to only include genes with ≥5 FPKM in at least 50% samples within a cancer type. This reduced the total number of genes from 19,801 to 13,250.

Next, we carried out clustering using the t-SNE unsupervised non-linear dimensionality reduction technique to determine if there were distinct gene expression patterns associated with different cancer types. Distinct clusters of different cancers were observed for only 14 of the analysed cancer types (Figure 1A). Upon examination, we observed substantial variability and skewed data distributions due to the high expression levels of certain genes in a subset of cancer samples. The data were then log-transformed as recommended for such skewed data distribution [36]. This reduced data variance to 1.46 from 25881672.21 and restricted the expression values between 0 and 5.15 (Supplementary Table 1).

**Figure 1:**
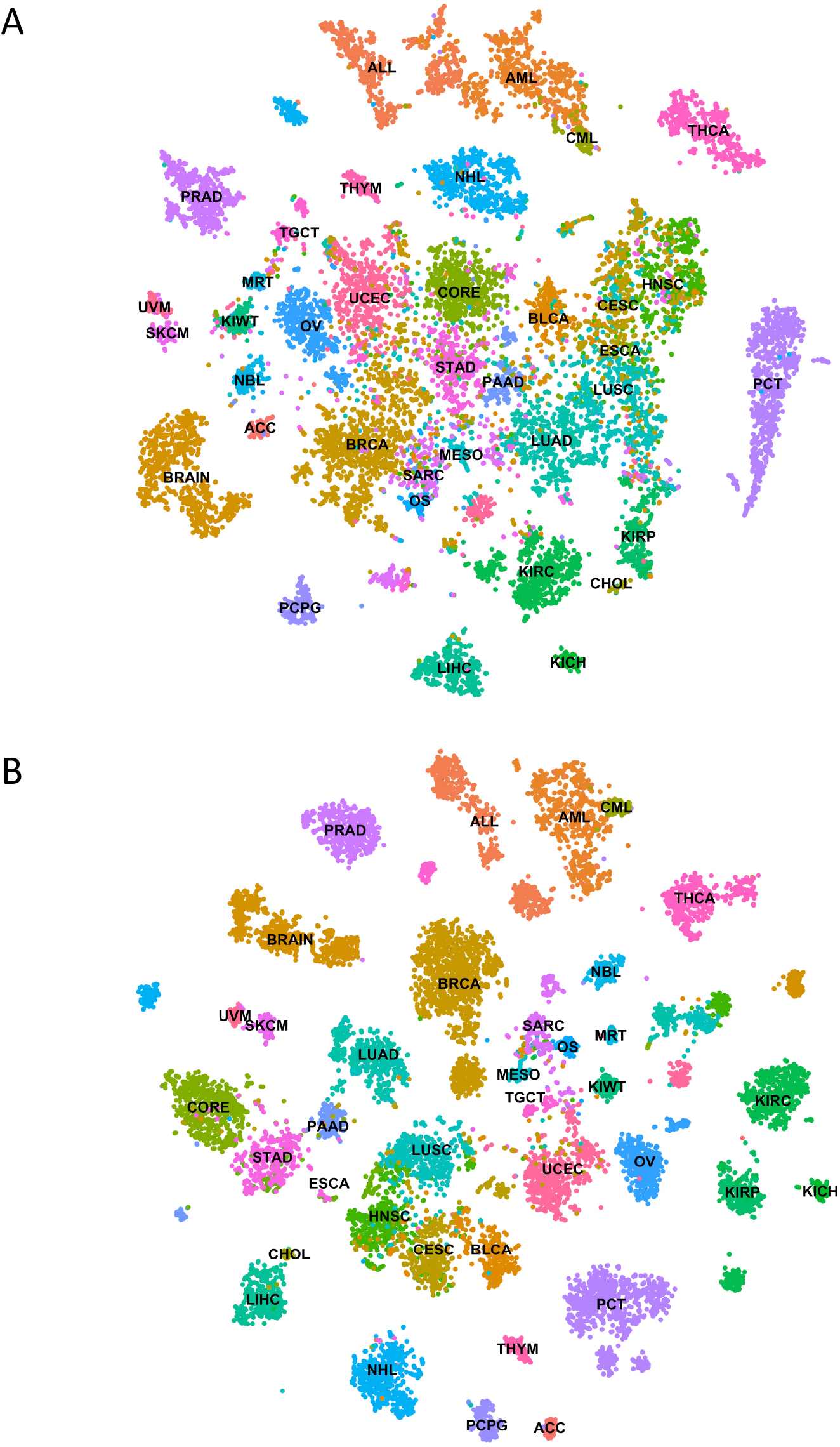
Unsupervised clustering of TCGA RNA-seq data using t-SNE. A) When FPKM values were used for the unsupervised clustering, did not separate cancer types into distinct groups. B) While the log10-transformed FPKM data exhibited better separation between cancer types

When log-transformed data was used for the t-SNE unsupervised clustering, it exhibited better separation between cancer types (Figure 1B). As expected, cancer types originating from the same tissues or cell types formed overlapping or proximal clusters due to similar underlying gene expression profiles, such as colon and rectum (CORE), skin cutaneous melanoma (SKCM) and uveal melanoma (UVM), acute myeloid leukemia (AML) and chronic myeloid leukemia (CML), uterine carcinosarcoma and uterine corpus endometrial carcinoma (UCEC) (Figure 1B). Liver hepatocellular carcinoma (LIHC) and cholangiocarcinoma (CHOL) are also clustered near one another owing to their proximity. Interestingly, cancers originating from squamous cells of different tissues such as lung squamous cell carcinoma (LUSC), head and neck squamous cell carcinoma (HNSC), cervical squamous cell carcinoma and endocervical adenocarcinoma (CESC), and bladder urothelial carcinoma (BLCA) were clustered near each other relative to adenocarcinomas of the same tissue types. Esophageal carcinoma (ESCA), which consisted of both adenocarcinoma and squamous cell carcinoma tissues, formed overlapping clusters with stomach adenocarcinoma (STAD) and squamous cell cancers. These clustering patterns reveal that unique gene expression profiles are associated with different cancer types.

### Development and training of a deep neural network model

We developed a pan-cancer classification model using deep neural networks (DNN) and RNA-seq gene expression data from TCGA. The choice of the network architecture for this model was data-driven. For example, while convolutional neural networks are preferred when data contains spatial patterns e.g. images. However, these spatial patterns may not be implicit in the gene expression matrix because differently arranged gene matrices will produce different spatial patterns. While recurrent neural networks (RNN) have recurrent blocks which act like a memory state that allow them to learn temporal information from the data [37]. Thus, they are preferred when input data is composed of sequential information with time dependency between values such as weather and speech data. Therefore, we opted for DNN based architecture as it is better at learning latent patterns in data where input variables are independent of one another. Our first DNN model was composed of 5 dense hidden layers with 50 nodes, and its accuracy was below 40%. The learning capacity of the model was improved by adjusting the number of hidden layers and nodes which resulted in improving model accuracy over 95%. Then we performed a grid search to find the best optimization algorithm and learning rate. For this, we used algorithms: Adam[30], Stochastic gradient descent[38], and RMSprop[39] optimizers with learning rates of 0.001, 0.001, and 0.00001. Our final DNN model consisted of 5 hidden layers with 500, 250, 125, 100, and 75 nodes, respectively (Figure 2). It was trained using the Adam optimization algorithm to minimize the categorical cross-entropy loss function, and the learning rate was set to 0.00001. The model architecture and training were implemented on an NVIDIA Tesla M60 GPU with 8GB RAM using TensorFlow 2.0 and Python3 [40]. It took 103 seconds to train the model. Figure 3 shows the accuracy and loss function during the training process of the model. Our DNN model took only 5 epochs to reach more than 90% accuracy and loss (categorical cross-entropy) as low as 0.28 without showing obvious overfitting. During training, two data splits were used; one for training and another for validation so as to monitor the model performance and to prevent any overfitting using early stop. The overall training accuracy of the model was 99.64%, and the validation accuracy was 97.40%. Even after repeated training of the model using randomly split data, we obtained consistent results (Figure S1).

**Figure 2:**
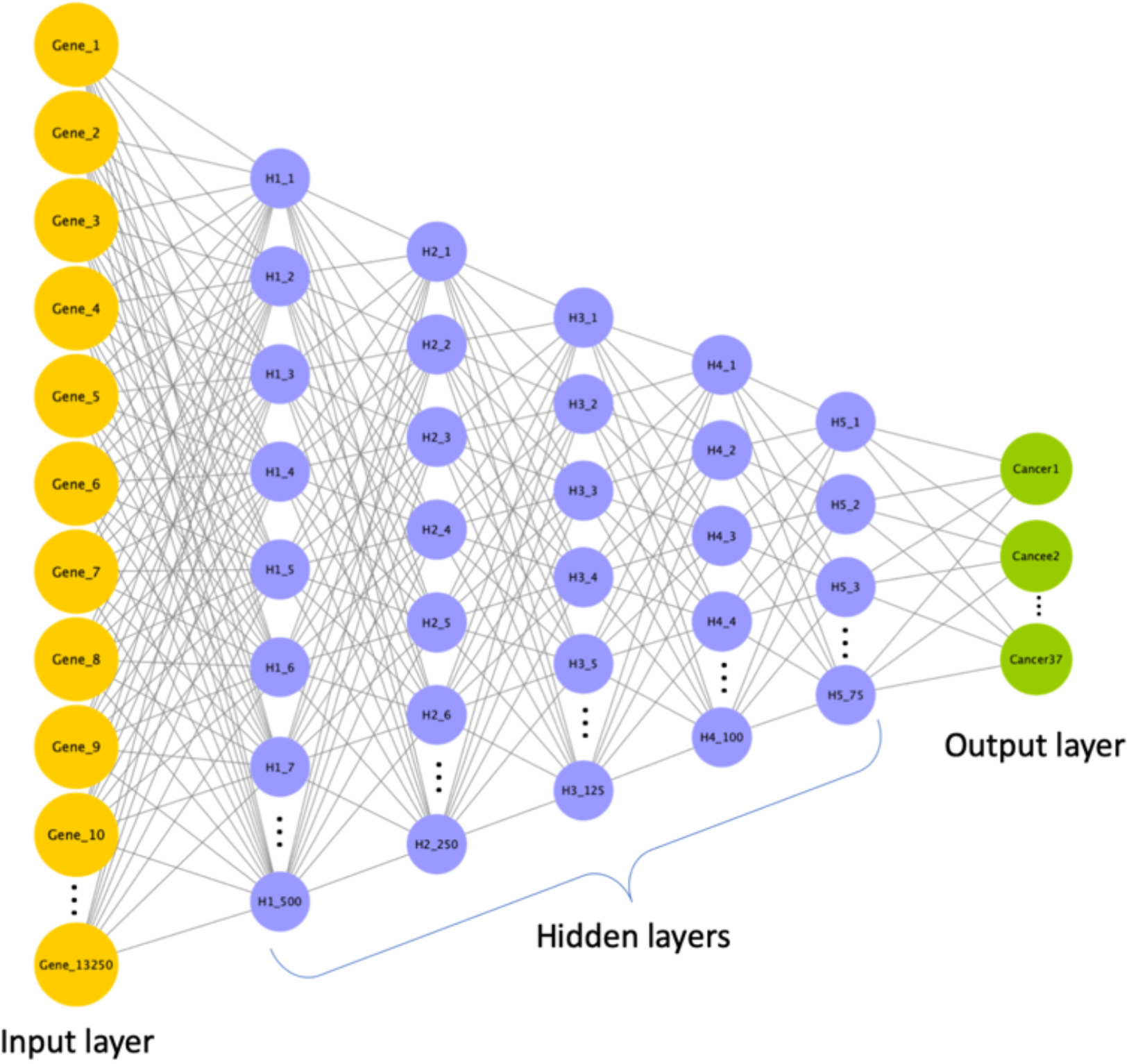
Architecture of deep neural network model. The DNN model consists of an input layer, five hidden layers and an output layer. The input layer takes input FPKM values for 13250 genes. Hidden layers perform nonlinear transformation using rectified linear unit activation function to distinguish between cancer types. There 5 hidden layers with 500, 250, 125, 100 and 75 nodes. Lastly, the output layer consists of 37 nodes each represent one of the cancers.

**Figure 3:**
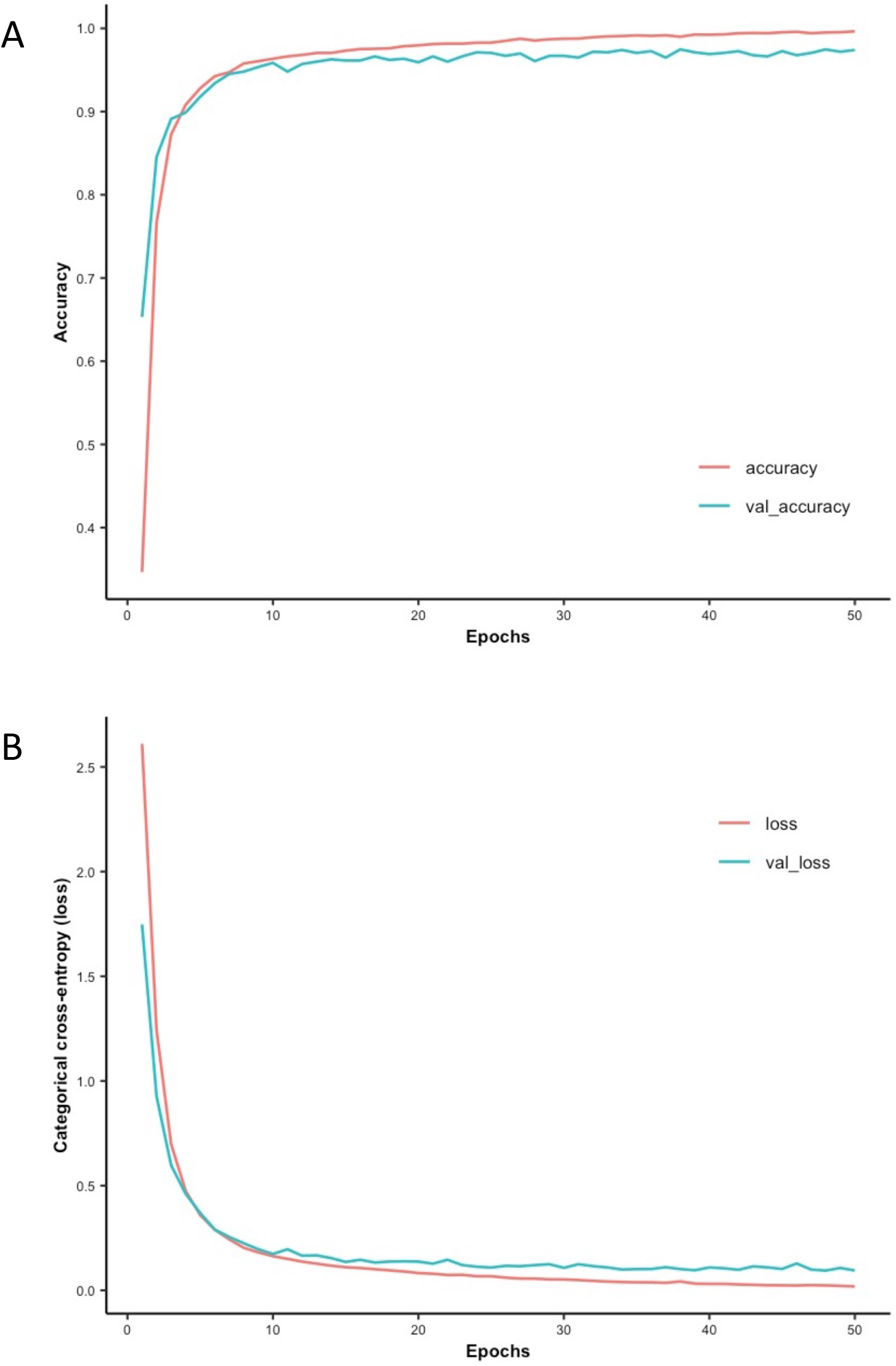
Accuracy and loss curve of the DNN model. A) The DNN model achieves 90% accuracy within five epochs on both training and validation datasets. B) The DNN model exhibits categorical cross-entropy (loss) that becomes negligible i.e. 0.0185 (training) and 0.0949 (validation).

### The trained DNN model accurately predicts cancer types based on gene expression profiles

To evaluate the performance of our DNN model, we next used it to classify a test dataset containing gene expression data from the 2,848 samples corresponding to 37 different cancer types (Table 1). The model accurately predicted the cancer type for 2,772 tumor samples, and misclassified 76 samples (Figure 4). The overall model accuracy on test data was 97.33%, which was slightly less than the training accuracy, which was 99.64%. The weighted averages of precision, recall, and f1-score values were 97.37%, 97.33%, and 97.33%, respectively. For most cancers, the test accuracy was above 95%, and it was 100% for the following 11 cancer types: adrenocortical carcinoma, kidney Wilms tumor, mesothelioma, neuroblastoma, ovarian serous cystadenocarcinoma, plasma cell tumors, prostate adenocarcinoma, skin cutaneous melanoma, testicular germ cell tumors, thyroid carcinoma, and uveal melanoma. The test accuracy for bladder urothelial carcinoma, esophageal carcinoma, kidney renal clear cell carcinoma, lung squamous cell carcinoma, malignant rhabdoid tumors, and osteosarcoma were between 90-95%. The test accuracy was below 90% for four cancer types: cholangiocarcinoma, chronic myeloid leukemia, kidney chromophobe, and stomach adenocarcinoma.

**Figure 4:**
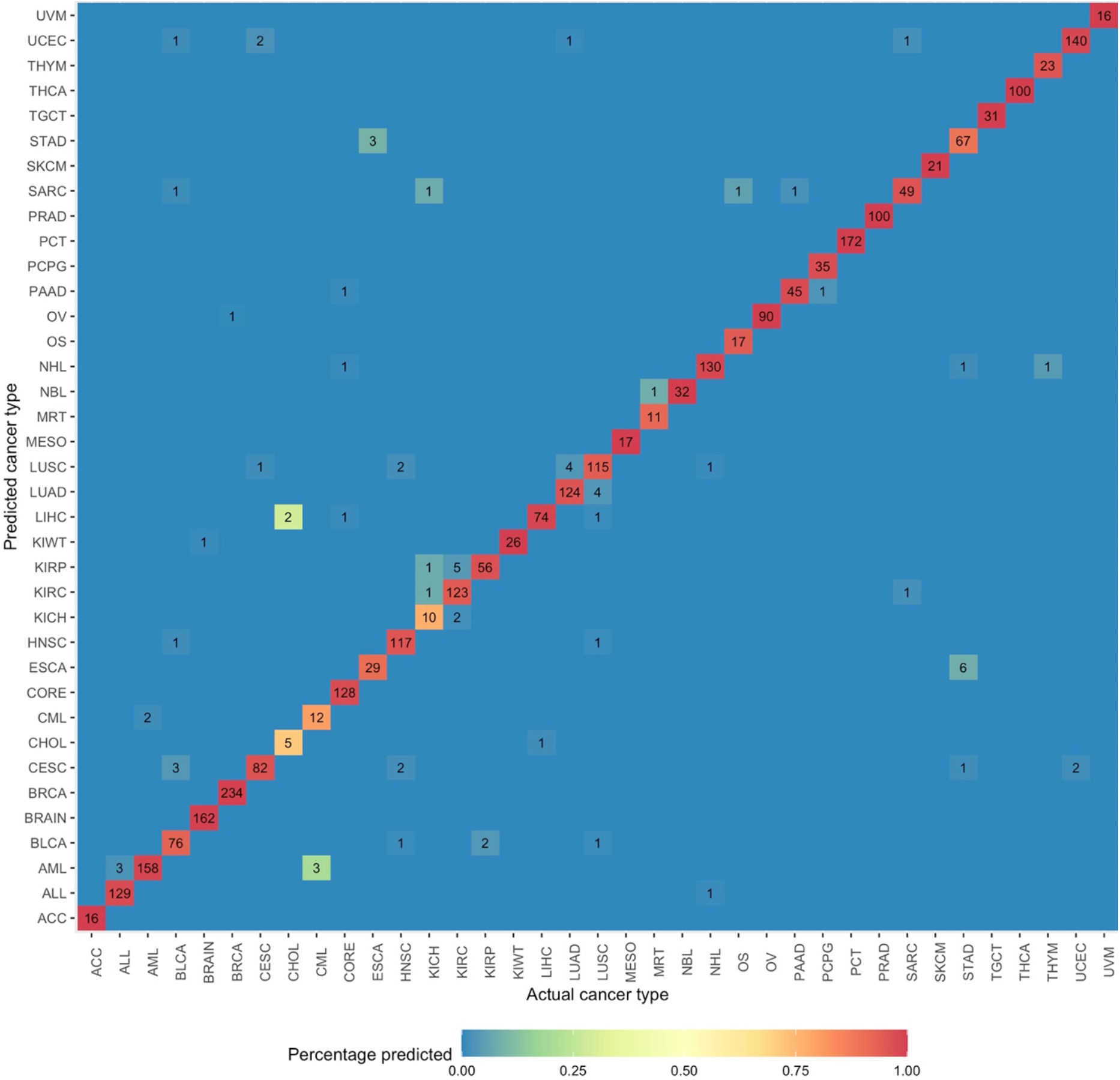
Performance of model on the test dataset. The majority (2772; 97.33%) of samples were accurately classified by the DNN model, as shown in this heatmap. However, there was some misclassification within particular organ systems including the kidneys and lung.

The model performed poorly on cholangiocarcinoma samples, with a test accuracy of 71.42%. This may be due to the limited number of samples (N=56) and the anatomical location of this cancer type. Small sample sizes hinder model learning and consequently result in the construction of an inaccurate classifier. Cholangiocarcinoma originates from the liver, and performing a biopsy of these tumors without liver cell contamination is non-trivial. Hence, cholangiocarcinoma samples often exhibit gene expression profiles similar to those associated with liver hepatocellular carcinoma. This was also evident from t-SNE unsupervised clustering results wherein these two cancer types formed proximal clusters (Figure 1B). Even though only 2 (out of 7) samples were misclassified, it significantly reduced test accuracy for cholangiocarcinoma. Interestingly, both misclassified samples were predicted as liver hepatocellular carcinoma.

Classification ambiguities were also observed among different cancers that arise from the same organ systems. For example, three acute lymphoid leukemia samples were predicted as being acute myeloid leukemia samples, seven kidney renal clear cell carcinoma samples as kidney renal papillary cell carcinoma or kidney chromophobe samples, four Lung squamous cell carcinoma samples as Lung adenocarcinoma samples, four lung adenocarcinoma samples as lung squamous cell carcinoma samples, and two cervical squamous cell carcinoma and endocervical adenocarcinoma samples as uterine corpus endometrial carcinoma samples. Similarly, six stomach adenocarcinoma samples were predicted as being esophageal carcinoma samples, and three esophageal carcinoma samples were classified as stomach adenocarcinoma samples. Classification ambiguity among cancers originating from proximal tissues also highlights the challenges associated with their diagnosis. For example, biopsy sites for 70% of esophageal carcinoma samples were in the lower third of the esophagus closer to the gastroesophageal junction. The precision score was lowest for esophageal carcinoma (82.85%), as six stomach adenocarcinoma samples were predicted as being esophageal carcinoma samples. Cholangiocarcinoma cancer had the lowest recall (sensitivity) score and f1-score values at 71.42% and 76.92%, respectively.

We performed five-fold cross-validation of our model by randomly dividing TCGA data into 5 datasets. Each time, we used one of those datasets as a test set and the remaining four as a training set for the model. Our model achieved consistent results in this cross-validation analysis, with a training accuracy of more than 99% (Table S1). Precision, recall, and f1-scores were also consistent. Overall, the model predicted different cancer types with over 97% accuracy.

### Identification of cancer-specific gene expression signatures

We next used SHAP values to interpret our model and to identify the gene signatures therein [31]. SHAP is a method that scores for each feature according to its contribution to the results of the model. A feature with the highest SHAP value is the major contributor to the output of a model [31]. To identify gene signatures, we developed 10 DNN models with the same architecture. To ensure that non-identical data was used during model construction, for each model, the TCGA data was randomly divided into training (70%), validation (10%), and test (20%) datasets. The performance of all 10 models was consistent and they all achieved 99% and 97% accuracy during training and testing, respectively (Figure S1). Next, we calculated SHAP values for each gene using the test dataset of the respective model. In each model, the top 20 genes were selected for each cancer type based on absolute mean SHAP values. Then, we calculated the number of times a gene was among the top 20 in a particular cancer type, henceforth referred to as occurrence frequency. Only genes with an occurrence frequency of at least 5 for a given cancer type were selected. This resulted in the selection of 507 candidate gene signatures. Subsequently, the fold change was calculated for each gene by comparing its median FKPM value across cancer types. Genes enriched with log2(fold change) ≥1 in not more than in 5 cancer types were selected as gene expression signatures. As a result, we identified 448 gene expression signatures. Similarly, we repeated the above procedure to identify additional gene expression signatures omitting these 448 already identified signatures. During this second iteration, we identified 310 additional gene expression signatures, as well as 218 in the third iteration. We stopped at the third iteration because we were unable to detect any specific gene signature for bladder urothelial carcinoma, esophageal carcinoma, mesothelioma, and sarcoma. In total, we identified 976 gene signatures across 37 cancer types (Figure 5 and supplementary table 2). Of these, 636 (65.16%) signatures had an occurrence frequency of 7 or above, while 550 (56.35%) signatures were specifically enriched in only one cancer type and 43 (4.4%) gene signatures were found to be enriched in 5 different cancer types. There were 55 genes that exhibited liver cancer-specific expression patterns. Signatures with an occurrence frequency of 10 that were enriched in only one cancer type can be considered to represent a highly specific biomarker. There were 134 such gene signatures associated with 27 different cancer types in the present study (Supplementary Table 2). These 134 genes included well-known cancer biomarkers such as KLK3, which is specific to prostate cancer, thus recapitulating existing biomarker results. Other examples included GFAP in brain cancer, CRYGN in Thyroid carcinoma, and NOX1 in colorectal cancer (Figure 6). The successful detection of these cancer-specific gene signatures demonstrates the validity of our approach.

**Figure 5:**
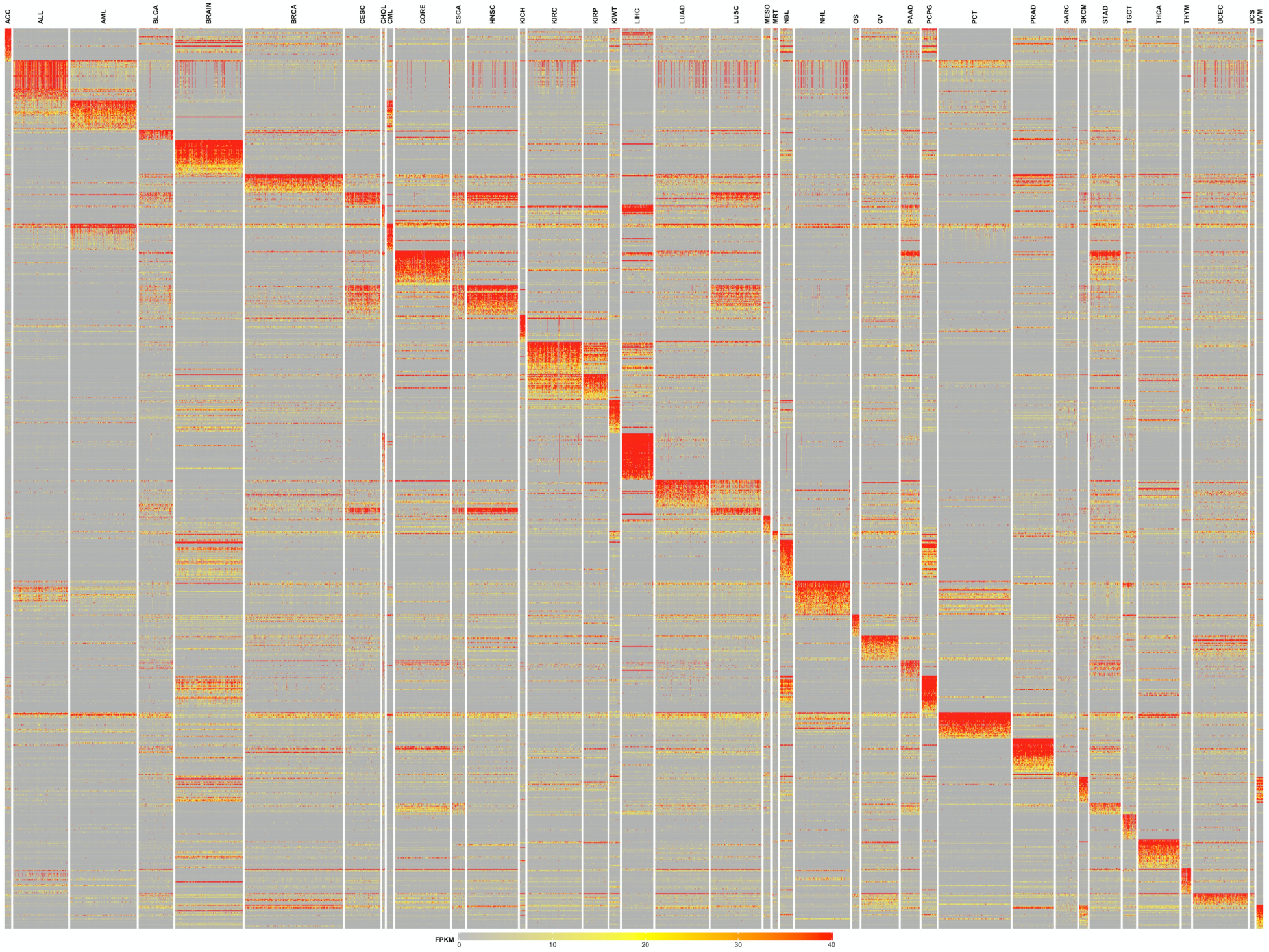
Gene signatures exhibit cancer-specific enrichment. Heatmaps demonstrating gene expression signatures across the TCGA data, revealing that many signatures exhibited cancer-specific expression patterns with the exception of sarcoma.

**Figure 6:**
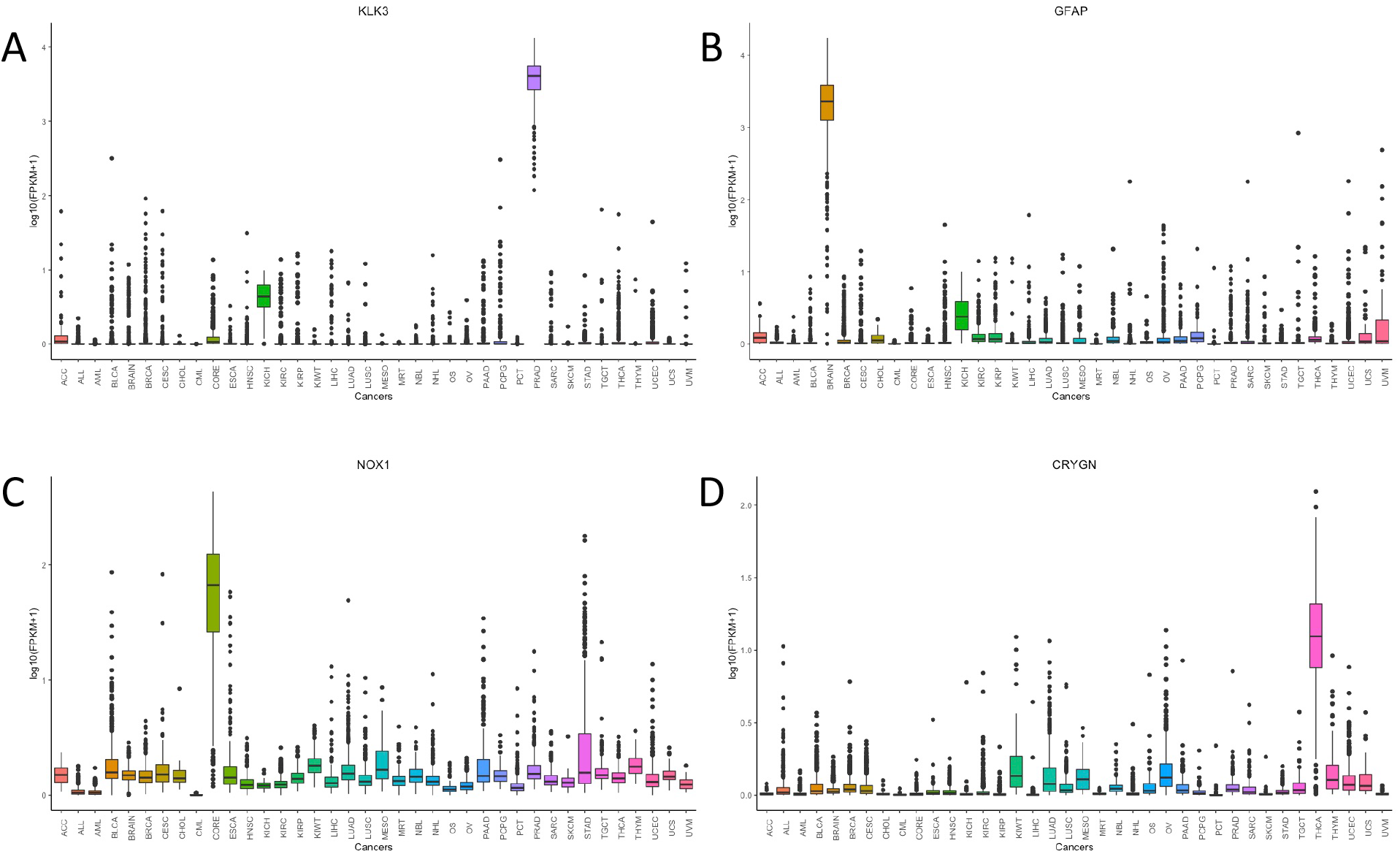
Genes signatures exhibit cancer-specific expression. Box plots showing expression of A) KLK3, B) GFAP, C) NOX1 and C) CRYGN across 37 cancer types in TCGA data. Median FPKM values of these genes were significantly higher in their respective cancer compared other cancers.

To evaluate the validity of gene signatures identified through our model interpretation, we built a new DNN model using the expression profiles of only these 976 gene signatures. The model architecture remained the same with the exception of the input layer, which was reduced from 13,250 to 976. The model exhibited consistent performance comparable to the original model and achieved 99% and 97% training and testing accuracy, respectively. Similarly, a weighted average of precision, recall, and f1-score values was consistent with the original model (97%). The model achieved consistent performance even during five-fold cross-validation (Figure S2 and Supplementary Table 3). We performed unsupervised t-SNE clustering of complete TCGA data by supplying expression values corresponding to the 976 included gene signatures. It produced distinct clusters for the majority of cancers (Figure S3). As expected, related cancers formed proximal clusters. The melanoma subtypes SKCM and UVM formed adjacent but well-separated clusters. These results demonstrate that our DNN model captures cancer-specific gene expression signatures through enhanced pattern recognition.

### Biomarkers for highly prevalent cancer types

We compared the cancer-specific gene expression signatures identified in our study with the list of United States Food and Drug Administration (FDA)-approved gene expression-based *in vitro* cancer diagnostic tests (https://www.fda.gov/medical-devices/vitro-diagnostics/nucleic-acid-based-tests). There are 10 gene expression-based tests available for five cancers including breast cancer, chronic myeloid leukemia, colorectal cancer, and prostate cancer (Table 2). The prostate cancer test is based on the expression of the PCA3 gene, which is an antisense RNA. As our DNN model was only based on protein-coding genes, it was excluded from the comparison. Discontinued gene symbols were also excluded. Only 12 out of 976 gene signatures overlapped with genes included in FDA-approved tests (Supplementary Table 4). All of these were associated with FDA-approved breast cancer tests produced by Veridex, LLC (1 gene), Agendia BV (3 genes), and Nanostring Technologies (8 genes). However, only 2 of those 12 genes (SCGB2A2 and SCUBE2) overlapped with our breast cancer signatures. The remaining 10 genes were assigned to cancers other than breast cancer in the present study. For example, the MLPH and MIA genes were identified as components of signatures associated with UVM and SKCM, respectively. Both of these genes were overexpressed in UVM and SKCM, respectively, relative to other cancers (Figure 7A and 7B). Thus, it is more appropriate to use these genes for UVM and SKCM diagnosis. Furthermore, 8 of the Agendia BV panel genes and 1 gene from the Nanostring technology panel did not meet our minimum expression threshold of 5 FPKM (Supplementary Table: 2). When these genes used in these FDA-approved tests of breast cancer tests were compared across cancers using TCGA data, a majority of the genes were detected in many cancer types and had no specificity for any particular cancer types (Figure S3).

**Table 2:** FDA-approved cancers test based on tumor gene expression profiles.

|  | Caner type | Manufacturer | Use | Total genes |
| --- | --- | --- | --- | --- |
| 1 | Breast Cancer | Nanostring Technologies | Prognostic | 50 |
| 2 | Breast Cancer | Agendia BV | diagnostic (metastasis) | 66 |
| 3 | Breast Cancer | Veridex, LLC. | diagnostic (metastasis) | 2 |
| 4 | Breast Cancer | BIOGENEX LABORATORIES, INC. | diagnostic | 1 |
| 5 | Chronic Myeloid Leukemia | ASURAGEN, INC. | diagnostic | 2 |
| 6 | Chronic Myeloid Leukemia | Cepheid | diagnostic | 2 |
| 7 | Chronic Myeloid Leukemia | Bio-Rad Laboratories, Inc. | diagnostic | 2 |
| 8 | Chronic Myeloid Leukemia | MolecularMD Corporation | diagnostic | 2 |
| 9 | Colorectal Cancer | Ventana Medical Systems | diagnostic* | 5 |
| 10 | Prostate Cancer | Gen-Probe, Inc. | diagnostic | 1 |

**Figure 7:**
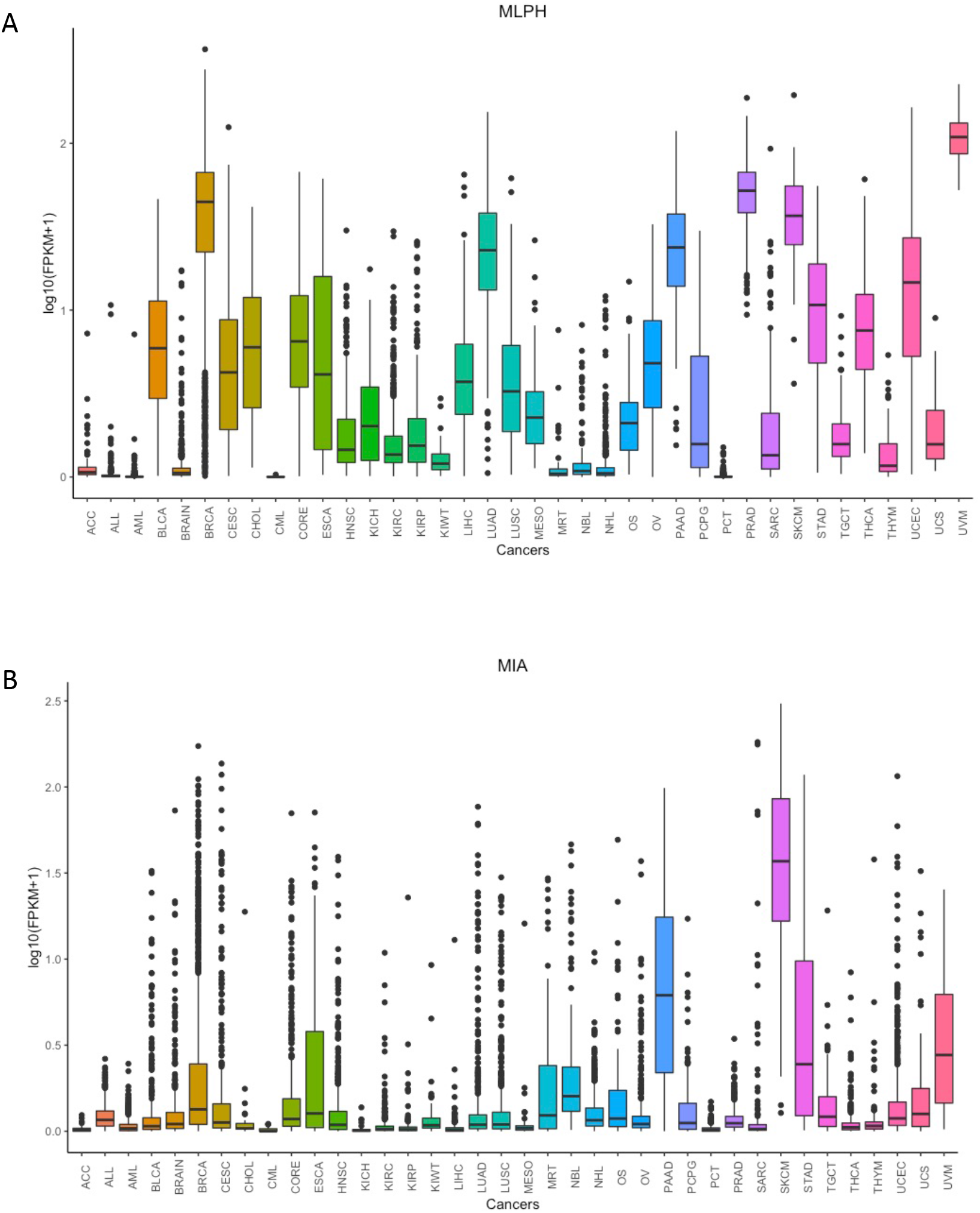
MLPH and MIA genes exhibit higher expression in melanoma cancers (SKCM and UVM) relative to breast cancer. Box plots showing expression A) MLPH and B) MIA genes across all cancers. Both genes had higher enrichment in melanoma cancers (SKM and UVM) compared to breast cancer (BRCA).

### Keratins for cancer diagnosis

Most epithelial cells express keratins, and the human genome encodes 54 different keratins, some of which exhibit cell- or tissue-specific expression patterns [41]. Owing to their diverse expression patterns, keratins are widely used in the histopathological evaluation of cancers [42]. For example, keratins KRT8, KRT18, and KRT19 are expressed in most adenocarcinomas [41]. Ovarian cancer cells are KRT 7^+^/ KRT 20^−^, whereas prostate cancer cells are negative for both KRT7 and KRT20 [43, 44]. The use of keratins as biomarkers of particular cancer types was common prior to the availability of pan-cancer gene expression datasets, but it is now possible to evaluate tissue-specific keratin expression patterns in order to determine whether they are the most specific biomarkers for a particular tumor type. Here, we compared the keratins used in the diagnosis of most common cancers such as breast cancer, prostate cancer, colorectal cancer, and skin cancer with our gene expression signatures. Surprisingly, our findings contradicted some published observations. For example, previous studies have reported that KRT8, KRT18, and KRT19 are expressed in most cancers including skin cancer [42], whereas our analysis revealed that they were depleted in skin cancer samples (Figure 8). Similarly, expression of KRT5, KRT6, KRT14, and KRT17, which are known to be expressed by both breast and skin cancer, was inconsistent with the expression patterns observed in the TCGA database. These keratins exhibited significantly low expression levels in almost half of the assessed skin cancer samples (Figure 8). Interestingly, KRT10, which is considered to be a skin cancer specific keratin, was detected in all cancers including brain and blood cancers. Despite their widespread use in histopathological contexts, our data suggest that there may be better alternatives to keratins when assessing particular cancer types, given that our gene expression signatures exhibited better tissue specificity than did keratins (Figure 5). Some of the skin cutaneous melanoma (SKCM) signatures were expressed in uveal melanoma (UVM) samples, but this was considered to be acceptable given that both UVM and SKCM are subtypes of melanoma. Similarly, some of the colorectal signatures were associated with stomach samples, albeit with variable expression levels. This is likely because these two cancers are both forms of gastrointestinal malignancies. As such, our pan-cancer gene expression signature may be of value for the development of novel immunohistochemical markers for cancer diagnosis.

**Figure 8:**
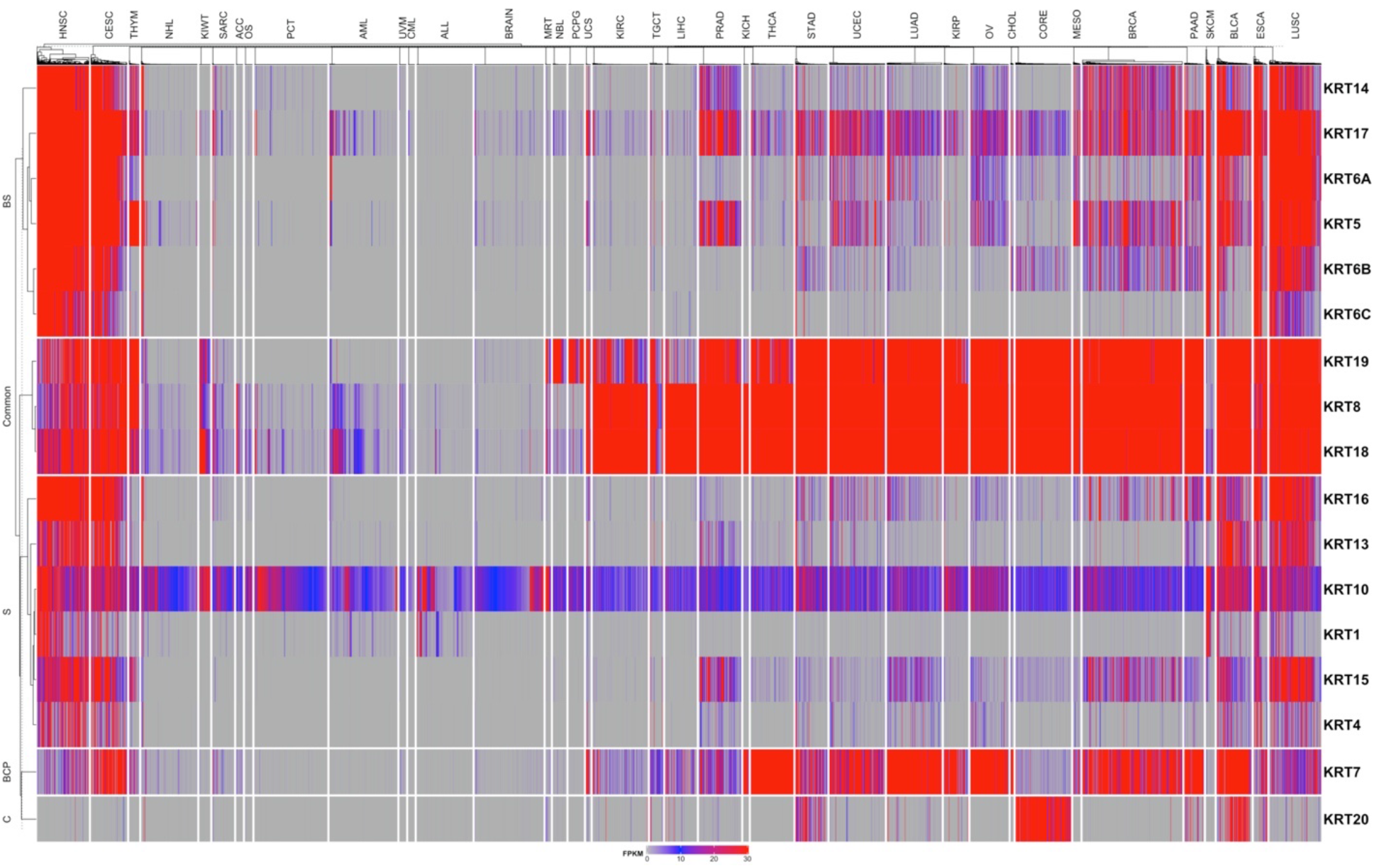
Heatmap showing expression of keratins used in diagnosis of BRCA, PRAD, CORE, and SKCM. As it can be seen from the heatmap that keratins used in the diagnosis cancer do not show cancer-specific expression.

## Discussion

One of the primary goals of cancer genomics and transcriptomics studies is the identification of cancer biomarkers and therapeutic targets. While many studies have been conducted in an effort to achieve this goal, the majority have focused on specific cancer types without utilizing molecular data from other cancers for comparison. The availability of cancer genome sequencing and transcriptome sequencing data from multiple cancer types has enabled pan-cancer analyses aimed at identifying cancer-specific biomarkers and therapeutic targets. In this study, we utilized transcriptomic data from 14,237 cancer samples in TCGA to carry out pan-cancer analyses identifying gene expression signatures associated with different cancer types. We used a deep learning approach to conduct pan-cancer classification and to define associated gene expression signatures. Our model is capable of identifying particular cancer types with greater than 97% accuracy based upon gene expression data alone. Classification ambiguities were primarily observed when comparing different cancer subtypes within a given organ system such as tumors of the blood, kidney, gastrointestinal organs, and lung. This is likely attributable to the high degree of molecular similarity between different cancer types originating from the same tissues. These results may also provide therapeutic opportunities for the repurposing of cancer drugs. In this study, we successfully implemented a SHAP method and identified a comprehensive list consisting of 976 pan-cancer gene expression signatures. These gene signatures were evaluated for their ability to accurately identify different cancer types. In order to do this, we developed an independent DNN model for which the inputs were the expression values of these 976 genes across different cancer types. The model showed consistent performance, achieving over 97% accuracy on an independent test dataset. Statistical measures that are not biased towards over-represented cancer types including precision, recall, and f1-score values were also above 97%, and the five-fold cross-validation of our model yielded consistent results.

Our analysis resulted in the identification of a number of cancer biomarkers across different cancer types. These included known markers confirming the validity of our approach. In addition, we identified several novel biomarkers that exhibited cancer-specific expression patterns. Such cancer-specific gene expression signatures may offer value for the development of gene panels that can be used to screen and diagnose cancers. Secreted cancer-specific marker proteins can also be leveraged to monitor disease progression and therapeutic responses. Circulating tumor cells (CTCs) similarly offer value for disease diagnosis and monitoring. They are also useful to determine appropriate therapy. However, the numbers of CTCs in patient blood are very low relative to the number of other blood cells. Extant methods isolating CTCs are non-specific and inefficient, while cancer-specific surface markers can be of great value for such isolation. The model and gene signatures identified in this study can also be leveraged to establish the tissue of origin in patients suffering from cancers of unknown primary, which is critical to providing appropriate treatment aimed at improving patient outcomes. Cancer of unknown primary account for 2-5% of all cancer diagnoses, and represent a major area of unmet clinical need. Cancer-specific biomarkers can also be used in imaging contexts. The targeted treatment of cancers is predicated on the identification of particular proteins on which cancers depend for their survival. Our list of cancer-specific gene signatures can therefore be used to guide the elucidation of novel therapeutic targets.

There are some limitations associated with our pan-cancer analysis-based assessment of cancer-specific gene expression signatures. For example, we have only considered genes that are expressed at sufficiently high levels (≥5 FPKM) in at least 50% of samples within a cancer type. While this will limit the number of false positive results, it may result in false negatives resulting in the loss of some markers that are expressed at low levels. We also focused only on genes that were overexpressed. Certain markers may be downregulated in specific cancer types, but they were not the focus of this study. Deep learning models aim to minimize the objective function, as a result, the model may not select certain genes that exhibit cancer-specific expression patterns if it has already identified many candidate genes that yield better signals associated with that cancer type which might help them achieve the object or minimize the loss. For example, NAT1 was expressed at higher levels in breast cancer compared to other cancer types, but the model did not select it.

In summary, we have developed a deep learning model that can accurately identify cancer types based on gene expression data. The cancer-specific gene expression signatures identified through this pan-cancer analysis may represent valuable biomarkers and therapeutic targets that can guide patient diagnosis and care.

## Notes

### Competing Interest Statement

The authors have declared no competing interest.

